# Hemi Manganese Exchangers 1 and 2 enable manganese import at the plasma membrane in cyanobacteria

**DOI:** 10.1101/2023.02.16.528846

**Authors:** Mara Reis, Fabian Brandenburg, Michael Knopp, Samantha Flachbart, Andrea Bräutigam, Sabine Metzger, Sven B. Gould, Marion Eisenhut

## Abstract

Manganese (Mn) is key to oxygenic photosynthesis as it catalyzes the splitting of water in photosystem II and functions as cofactor of multiple enzymes. A single ABC-type transporter, MntCAB, is so far established for the uptake of the metal under limiting conditions in cyanobacteria. It is unknown how Mn is imported under replete conditions. We identified two proteins in the cyanobacterium *Synechocystis* sp. PCC 6803, which are homologous to the unknown protein family 0016 (UPF0016) member manganese exporter (Mnx). In contrast to Mnx, which consists of six transmembrane domains, the new candidate proteins contain three transmembrane domains. Hence, we named them hemi manganese exchangers (Hmx) 1 and 2. Knock-out mutants in *hmx1* and/or *hmx2* showed sensitivity toward low Mn supplementation, and reduced intracellular Mn pools. Additional deletion of *mntC* hindered the cells to thrive unless the medium was supplemented with Mn to compensate for the depletion of their intracellular Mn pool. In accordance with the observed localization of Hmx1 and Hmx2 in the plasma membrane, we postulate a Mn uptake function for heteromeric Hmx1/2 across the plasma membrane under a wide range of Mn concentrations and a supporting role for the MntCAB system under Mn-limiting conditions. On the basis of their phylogenies, we propose that Hmx1 and Hmx2 are the ancestral progenitors of eukaryote-type UPF0016 proteins with six transmembrane domains. The Mn transport function of Hmx1/2 underscores this as a fundamental and ancient feature of the UPF0016 family. Potentially, Hmx1 and Hmx2 coevolved with the internalization of the oxygen-evolving complex.

## INTRODUCTION

The transition metal manganese (Mn) is of crucial importance across all kingdoms of life. Mn associates with proteins, metabolites, and nucleic acids (reviewed in Bosma et al., 2021), fulfills an activating role for several proteins, and serves as a direct cofactor of a variety of enzymes (Hänsch & Mendel, 2009; Schmidt & Husted, 2019). Mn-dependent superoxide dismutase is central in scavenging of reactive oxygen species, and also Mn^2+^-ions associated with uncharacterized small molecules are suggested to defend against oxidative stress (Jakubovics & Jenkinson, 2001; Marschner, 2012). Diverse enzymes of carbon metabolism (Marschner, 2012), glucosyltransferases (Breton et al., 2006), oxalate decarboxylase (Tottey et al., 2008), and enzymes involved in isoprenoid and amino acid biosynthesis are strictly Mn-requiring (Schmidt et al., 2020). In oxygenic photosynthetic organisms, Mn is of even higher importance, since it functions as the inorganic catalyst in the oxidation of water. Together with oxygen and calcium atoms, Mn ions form the Mn_4_O_5_Ca cluster and catalyze the light-driven splitting of water molecules into electrons, protons, and molecular oxygen, as part of photosystem II (Nelson & Junge, 2015). Consequently, photoautotrophic cyanobacteria have a 100-fold higher demand for Mn than non-photosynthetic bacteria (Keren et al., 2002).

Critical for proper provision of the cell with Mn is a sufficient uptake of the metal from the environment into the cytoplasm, and further integration into the target proteins and molecules. In bacteria, two different types of Mn importers are known: the <u>A</u>TP-<u>b</u>inding <u>c</u>assette (ABC) type transporter and the <u>N</u>atural <u>R</u>esistance-<u>A</u>ssociated <u>M</u>acrophage <u>P</u>rotein (NRAMP) type transporter (Bosma et al., 2021). Though NRAMP proteins, such as MntH (Kehres et al., 2000), are encoded by cyanobacterial genomes, their specific relevance in metal transport has not been investigated so far. In contrast, the ABC-type Mn transporter MntCAB (Bartsevich & Pakrasi, 1995; Bartsevich a& Pakrasi, 1996) is well studied in the model cyanobacterium *Synechocystis* sp. PCC 6803 (*Synechocystis*). Expression of the *mntCAB*-operon occurs under the control of the two-component system ManS/ManR (Ogawa et al., 2002; Yamaguchi et al., 2002). Sensing of Mn limitation induces the expression and installation of the high-affinity transporter in the plasma membrane and enables cell growth under such conditions (Bartsevich & Pakrasi, 1995). Intriguingly, a deletion mutant in *mntC* was still able to accumulate Mn inside the cells when (sub)micromolar amounts of Mn were present in the growth medium. This hinted at the presence of a second, yet unidentified Mn uptake system in *Synechocystis* (Bartsevich a& Pakrasi, 1996).

Another class of Mn transporter occurring in cyanobacteria and also eukaryotes, was only recently identified. The cyanobacterial founding member is the <u>Mn</u> e<u>x</u>porter (Mnx) (Brandenburg et al., 2017b), also named SynPAM71 (Gandini et al., 2017). Mnx resides in the thylakoid membrane and shuttles Mn from the cytoplasm into the thylakoid lumen to (i) assist in Mn provision to PSII and (ii) prevent detrimental overaccumulation of Mn in the cytoplasm (Brandenburg et al., 2017b). Mnx belongs to the <u>U</u>ncharacterized <u>P</u>rotein <u>F</u>amily (UPF) 0016. Most members of the family consist of two repetitions of a domain that contains a conserved ExGD motif in the first of three predicted transmembrane domains (TMDs), making them 6-TMD proteins (Demaegd et al., 2014). Transport of Mn was demonstrated for the UPF0016 members PAM71 (Schneider et al., 2016), CMT1 (Eisenhut et al., 2018; Zhang et al., 2018), PML1 (Hoecker et al., 2020), PML2 (Hoecker et al., 2020), and PML3 (Yang et al., 2021; He et al., 2022) in *Arabidopsis thaliana* (Arabidopsis). In other analyses, also Ca was determined to be a transport substrate (Wang et al., 2016; Frank et al., 2019).

Here, we identified and characterized two additional members of the UPF0016 in *Synechocystis.* <u>H</u>emi <u>m</u>anganese e<u>x</u>changer (Hmx) 1 and 2 are each 3-TMD proteins and jointly function as a constitutive Mn importer in the plasma membrane of *Synechocystis*. They likely represent the ancestral state from which the 6-TMD proteins, such as Mnx or PAM71, which is the only version of UPF0016 proteins occurring in eukaryotes, evolved through gene fusion.

## MATERIAL AND METHODS

### *Synechocystis* Strains and Growth Conditions

The glucose-tolerant Japanese strain of *Synechocystis* sp. PCC 6803 obtained from Martin Hagemann (University of Rostock, Germany) served as wild type (WT). Axenic cultures were routinously grown in BG11 medium adjusted with 20 mM HEPES-KOH to pH 7.5 (Rippka et al., 1979) in a shaker at 30 °C and 200 rpm, illuminated with 100 μmol photons m^-2^ s^-1^ constant white light. Growth medium of the mutant lines was supplemented with appropriate antibiotics, 50 μg mL^-1^ kanamycin (Km), 20 μg mL^-1^ spectinomycin (Sp), or 12.5 μg mL^-1^ gentamycin (Gm).

### Generation of *Synechocystis* Knockout and Double-Knockout Lines

The Δ*hmx1* knockout mutant was generated by introduction of the plasmid pUC-4S including a Sp resistance cassette after HincII digestion into the NaeI restriction site of the PCR-amplified (primers FB69 and FB70; Table S1) open reading frame of *slr*1170. The cloning vector pJET1.2 (ThermoFisher) served as vector backbone. To generate the knockout construct for *hmx2,* the open reading frame for *ssr*1558 including upstream and downstream sequences was amplified using the primers ME368 and ME369 (Table S1). The Km resistance cassette derived from the plasmid pUC-4K after digestion with HincII was inserted into the SmaI restriction site of *hmx2*. The construct for generating a Δ*mntC* mutant was produced by amplification of the open reading frame *sll*1598 using the primers ME163 and ME164 (Table S1). The Sp resistance cassette from pUC-4S was introduced into the MscI site. All restriction enzymes were purchased from New England Biolabs, USA.

Transformation, selection on Sp- or Km-containing BG11 plates and segregation of independent clones was verified by PCR analysis as described in (Eisenhut et al., 2006). The Δ*hmx1*/Δ*hmx2,* and Δ*hmx2*/Δ*mntC* double knockout mutants were generated by transformation of single mutants with the desired knockout-construct, following the same protocol.

### Subcellular localization experiments

For the generation of CFP-fusion protein constructs, the CFP-coding sequence and Gm resistance cassette from a vector as described in (Heinz et al., 2016) was utilized. Using the primers FB105 and FB106 (Table S1) XhoI and NheI restriction sites were added by PCR to the *hmx1* open reading frame including its 800 bp upstream region. Similarly, the 800 bp downstream region of *hmx1* was PCR amplified with EcoRI restriction sites added to both ends (primers FB107 and FB108; Table S1). Restriction digest and T4 DNA Ligase (all NEB enzymes) were used to first clone the 3’-region of *hmx1* downstream of a CFP and Gm resistance cassette, before *hmx1* including the 5’-region of *hmx1* was cloned upstream of the *cfp* gene. A GSGSG peptide linker separates the gene of interest and *hmx1* to allow proper folding of both proteins. As vector backbone, pJET1.2 (ThermoFisher) was used. A *hmx2:cfp* fusion was generated in the same way using primers FB109, FB110 and FB111, FB112 (Table S1). *Synechocystis* Δ*hmx1* and Δ*hmx2* cells were transformed with the constructs as described above, using Gm for selection. The antibiotics Km and Sp used for selection of the knockout of *hmx1* and *hmx2*, respectively, were omitted from the medium to allow replacement of the knockout alleles by the expression alleles. Successful transformation and segregation was verified by PCR using primers FB69/FB70 (*hmx1:cfp*) or ME368/ME369 (*hmx2:cfp*). For primer sequences see Supplemental Table S1.

For imaging, the cells were immobilized on microscopic glass slides by a thin layer of solid BG11 medium (1:1 mixture of 2-fold concentrated BG11 medium, with 24 mM sodium thiosulfate added and 3% [w/v] bacto agar). A Leica TCS SP8 STED 3X microscope with a HC PL APO CS2 100x/1.40 OIL objective was used. An argon laser at 488 nm and 70 W output intensity was used for excitation. Emission was detected using Leica HyD hybrid detectors from 470-530 nm (CFP) and 660-700 nm (chlorophyll). Microscopy was performed at the Center of Advanced Imaging, Heinrich-Heine University Düsseldorf.

### Drop Tests

The effect of varying amounts of MnCl_2_ on the different lines was tested on solid BG11 medium. Cultures were grown until mid-log phase and 5 d starved for Mn by cultivating in BG11 medium without MnCl_2_ (BG11 -Mn). Then, 2 μL of culture with an OD_750_ of 0.25 and subsequent 1:10, 1:100, and 1:1000 dilutions were spotted onto agar plates (BG11, pH 7.5 with 24 mM sodium thiosulfate added; solidified with 1.5% [w/v] bacto agar). The plates were supplemented with MnCl_2_ as indicated (1x MnCl_2_ = 9 µM MnCl_2_) and did not contain antibiotics. Plates were incubated under continuous white-light illumination of 100 μmol photons m^-2^ s^-1^ at 30 °C for 5 d.

### ICP-MS Measurements

Cells were washed with EDTA (20 mM HEPES-KOH, pH 7.5, and 5 mM EDTA) (Keren et al., 2002) before and after pre-cultivation under Mn-limiting conditions (BG11 -Mn) for 5 d, to ensure similar intracellular Mn concentrations in all lines. Before the experiment, cells were adjusted to an OD_750_ of 0.8 and treated with MnCl_2_ concentrations as given in the respective experiments. Before the experiment and 4 h after MnCl_2_ treatment, samples were taken and washed as described in (Brandenburg et al., 2017a). In short, to determine the total content of Mn in a sample, cells were washed two times with ice-cold HEPES (20 mM HEPES-KOH, pH 7.5) to preserve the periplasmic Mn storage. Additionally, the samples were washed two times with 4 mL Milli-Q grade (18 MΩ cm) water before further processing. To release the periplasmic Mn pool, a second set of samples was washed initially two times with HEPES containing 5 mM EDTA and subsequentially washed two times with Milli-Q grade water. The washed samples were re-suspended afterwards in 0.4 mL 65% nitric acid and digested for 3 h at 70 °C. The digested samples were diluted to ∼4 % nitric acid with 6.5 mL Milli-Q grade water. Elemental composition of the samples was determined by ICP-MS (Agilent 7700) at the CEPLAS Plant Metabolism and Metabolomics Facility, University of Cologne. The cell numbers of the samples were estimated using a cell counter (Beckman Coulter Z2).

### Sequence Analysis and Identification of Candidate Genes

Proteins of the UPF0016 were identified using Pfam (http://pfam.xfam.org) (Finn et al., 2016) and BlastP (https://blast.ncbi.nlm.nih.gov/Blast.cgi) analysis (Altschul et al., 1990). DNA and protein sequences were obtained from the genome database CyanoBase (http://genome.microbedb.jp/cyanobase). Clustal Omega (https://www.ebi.ac.uk/Tools/msa/clustalo) was used for sequence alignment (Sievers et al., 2011).

### Expression Analysis

The coefficient of variation (standard deviation divided by the mean to account and correct for transcript abundance level) was calculated for all genes in *Synechocystis* sp. PCC 6803 (Wulf et al., 2024) by mapping 46 publicly available expression datasets (SRX10706739, SRX10706741, SRX10706742, SRX10706743, SRX10706745, SRX10706747, SRX10706750, SRX3580390, SRX5980004, SRX5980006, SRX5980011, SRX5980016, SRX5980017, SRX5980018, SRX5980022, SRX7098395, SRX8102890, SRX8102891, SRX8102892, SRX8102893, SRX8102894, SRX8102895, SRX8102896, SRX8102897, SRX8102898, SRX8102899, SRX8102900, SRX8102901, SRX8844415, SRX8844416, SRX8844417, SRX8844420, SRX8844421, SRX8844428, ERX3642241, ERX3642242, ERX3642243, ERX3642245, ERX3642247, ERX3642249, ERX3642250, ERX3642253, ERX3642254, SRX7873832, SRX7873837, SRX9234496) using kallisto version 0.44 in stranded mode onto the transcriptome of *Synechocystis* sp. PCC 6803 (accession number NC_000911, downloaded on January 03, 2023, from NCBI https://www.ncbi.nlm.nih.gov/nuccore/NC_000911). Transcript per million values were exported to Excel and mean and standard deviation calculated using AVERAGE and ST.DEVP.

### Phylogenetic Analysis

A database of 5,655 complete prokaryotic genomes of the RefSeq database (O’Leary et al., 2016) was search via diamond blastp (Buchfink et al., 2015) using the “--very-sensitive” option. Since Mnx, Hmx1, and Hmx2 share significant sequence identity, exact pairwise global alignments were produced to determine whether queries with multiple hit are a Hmx protein or the fusion Mnx protein (Rice et al., 2000). Homologs were visualized using a presence-and-absence matrix, color-coding their pairwise local identity. The same set of seed sequences was used to search for eukaryotic homologs within a database of 150 eukaryotes (Ku et al., 2015). All significant hits with a maximum e-value of 1x10^-10^ and at least 25% sequence identity were retained for further analysis (Table S2).

## RESULTS

### The genome of *Synechocystis* encodes two half-sized proteins with respect to UPF0016 homologs

Transporters of the Mnx family (UPF0016) are critical for proper Mn distribution within the cyanobacterial (Brandenburg et al., 2017b; Gandini et al., 2017) and plant cell (Schneider et al., 2016; Eisenhut et al., 2018; Zhang et al., 2018; Hoecker et al., 2020; Yang et al., 2021; He et al., 2022). While we identified two plastid-targeted Mnx proteins for all members of the green lineage (Schneider et al., 2016; Hoecker et al., 2017; Eisenhut et al., 2018), in cyanobacteria just a single homolog was found. Closer inspection of cyanobacterial genomes, however, revealed the existence of two additional genes encoding more distantly related UPF0016 proteins. These proteins are half-size versions of the canonical Mnx proteins and contain only one cluster consisting of three TMDs including the conserved ExGD motif, of which two in tandem are characteristic for the UPF0016 family (Figure 1A). Accordingly, we named these proteins <u>H</u>emi <u>m</u>anganese e<u>x</u>changer (Hmx) 1 and 2. In *Synechocystis,* Hmx1 is encoded by *slr*1170 and Hmx2 by *ssr*1558. A BlastP run shows that Slr1170 and Ssr1558 have a 35% and 31% sequence identity to *Synechocystis* Mnx, respectively. Furthermore, we found Hmx1/Hmx2 orthologues to be encoded by neighboring genes in the majority (90 %) of the 172 cyanobacterial species we analyzed, while in only a few (10%) species, such as in *Synechocystis*, they were dispersed across the genome (Table S3). In those cases, where *hmx1* and *hmx2* genes are organized as a transcriptional unit, we frequently detected a third gene being part of the operon (Figure 1B). They are members of the UPF0153, yet their function remains unknown. One exception is *psb28-2* of *Cyanothece* sp. ATCC 51142, which was identified upstream of the *slr*1170 orthologue. Psb28-2 is involved in PSII and chlorophyll biosynthesis (Dobáková et al., 2009; Nowaczyk et al., 2012).

**Figure 1:**
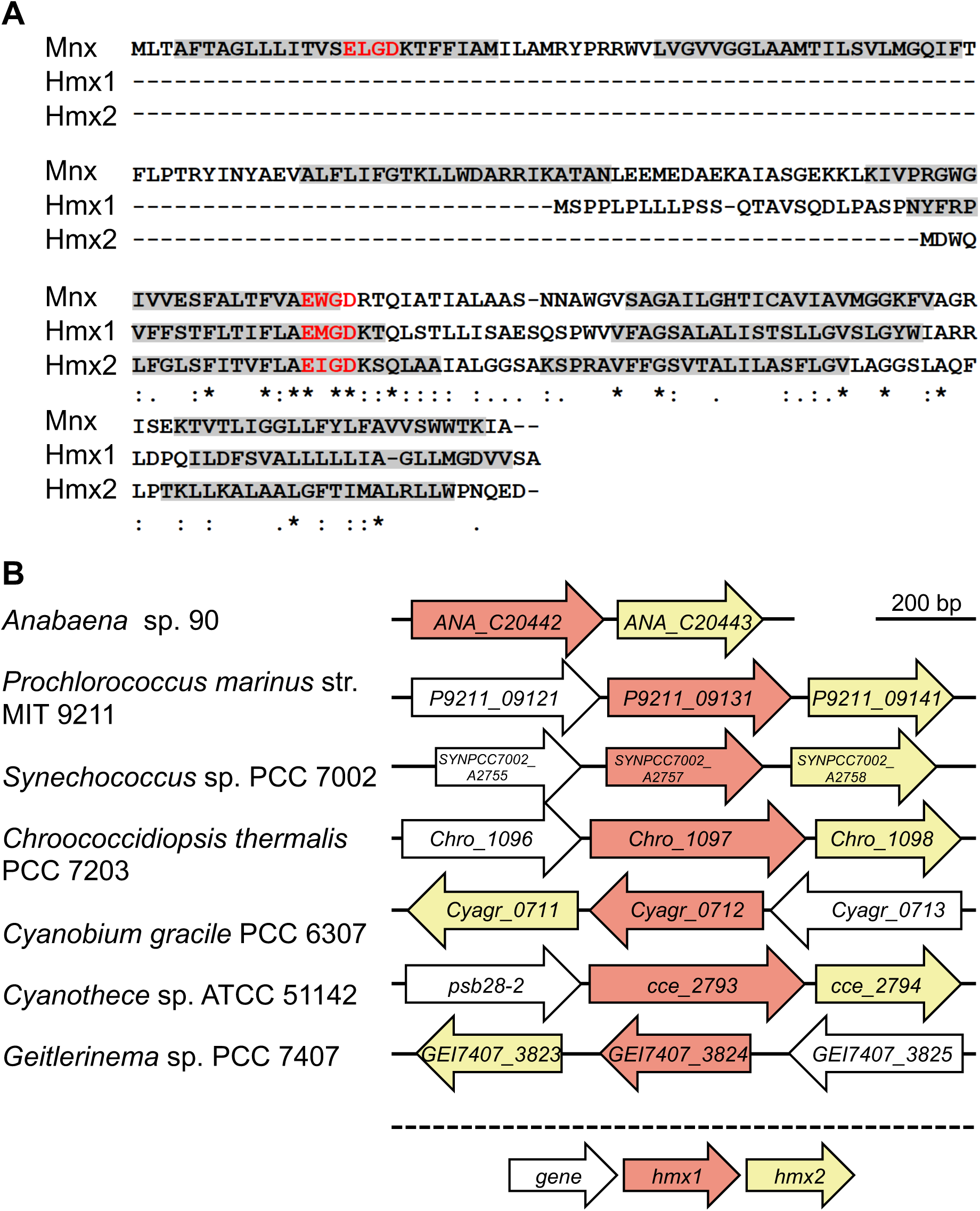
A) Amino acid alignment of UPF0016 members of *Synechocystis*. Grey boxes indicate predicted transmembrane domains. The conserved ExGD motif is highlighted in red. Asterisks (*) indicate strictly conserved residues and a colon (:) depicts residues with similar properties. B) Genomic organization of *hmx1* (orange) and *hmx2* (red) orthologs across some selected cyanobacteria. Additional genes as part of the transcriptional unit are depicted in grey. The direction of arrows indicates the orientation in the genome.

### Deletion of *hmx1* or *hmx2* impairs intracellular Mn accumulation

Due to the frequent genomic appearance of the genes in transcriptional units, we hypothesized that Hmx1 and Hmx2 interact and function in Mn transport, as other members of the Mnx family do. To test this hypothesis, single and double mutants of *hmx1* or/and *hmx2* were generated by insertional inactivation in *Synechocystis*. In the case of *hmx1,* a Sp resistance cassette and in the case of *hmx2,* a Km resistance cassette was inserted into the coding region leading to an interruption of the respective reading frame (Figure 2A). As verified by PCR, we obtained fully segregated mutant lines for Δ*hmx1,* Δ*hmx2,* and Δ*hmx1/*Δ*hmx2* (Figure 2B). All lines were tested for their susceptibility toward different Mn concentrations (Figure 2C). In contrast to the Δ*mnx* mutant, which is sensitive toward elevated Mn supply (Brandenburg et al., 2017b; Gandini et al., 2017), mutants in *hmx* genes were not susceptible to high but low Mn concentrations in the medium. Without any and with standard (1x, 9 µM MnCl_2_) Mn supply, single and double mutants displayed retarded growth. This impairment was improved by elevated (5x) Mn concentration in the medium. We also tested the mutantś sensitivity towards reduced supplementation with calcium (Ca) and iron (Fe) (Figure S1), since other known Mn transporters, such as members of the NRAMP, IRT, or the CAX family, are known to also use these substrates (Socha & Guerinot, 2014). While omitting Fe from the medium fully disabled growth of all strains, including that of the WT, depleting Ca only did not affect growth of any strain. Also, all other combinations did not result in different growth behavior of Δ*hmx1* and Δ*hmx2* mutants in comparison to the WT.

**Figure 2:**
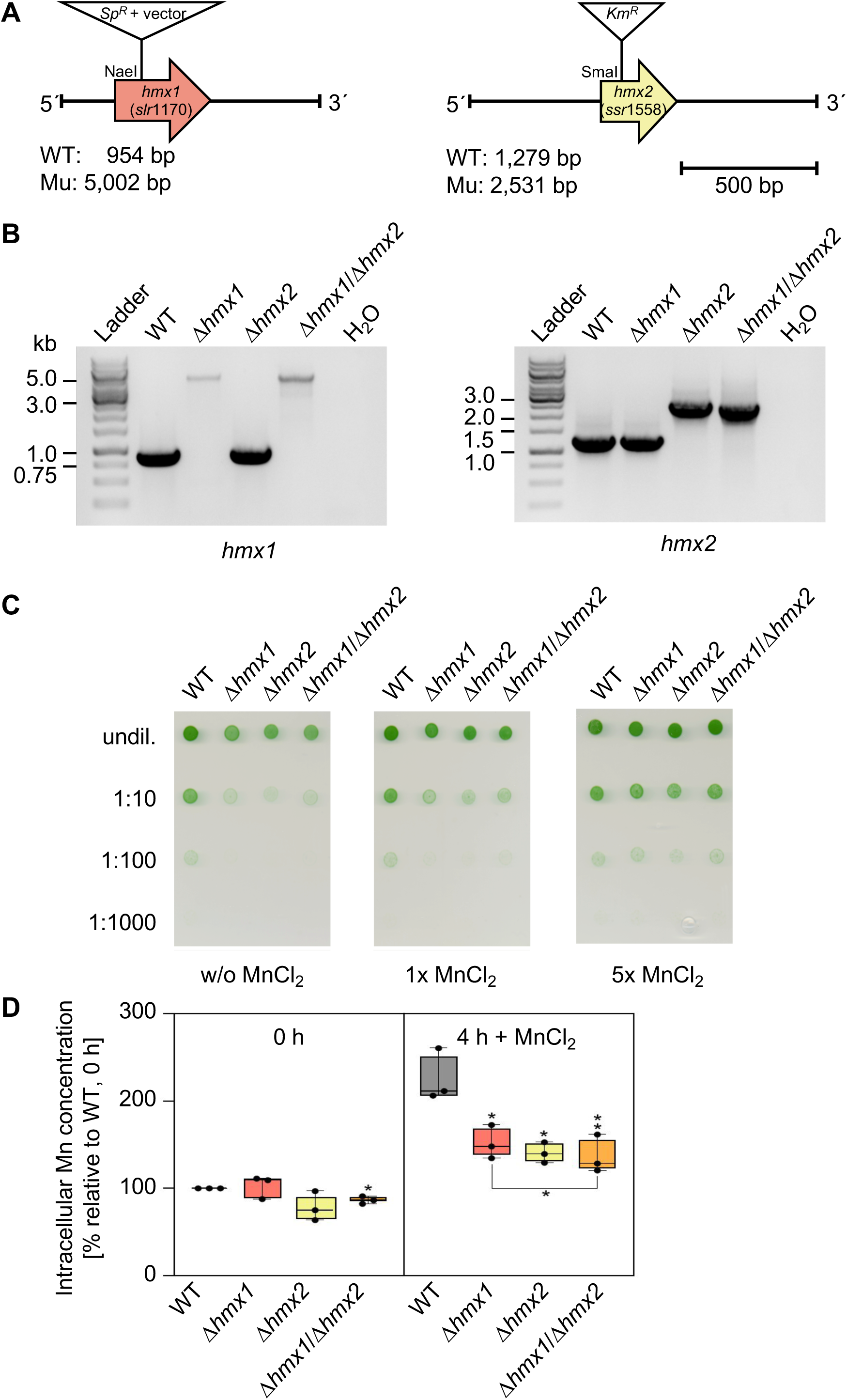
Analysis of Δ*hmx1* and Δ*hmx2* mutants. A) Scheme for the generation of mutants in *hmx1 (*Δ*hmx1)* and *hmx2* (Δ*hmx2*) by insertional inactivation. B) Verification of mutations in Δ*hmx1*, Δ*hmx2,* Δ*hmx1/*Δ*hmx2,* and Δ*hmx2/*Δ*hmx1* by genotyping. PCR analysis was performed with gDNA and gene specific primers *(hmx1*: primers FB69/FB70, WT = 954 bp; Mu = 5,002 bp. *hmx2*: primers ME368/ME369, WT = 1,279 bp; Mu = 2,531 bp). C) Droptest to monitor Mn sensitivity. Cells were washed with BG11 -Mn and adjusted to an OD_750_ of 0.25. Cells were diluted 1:10, 1:100, 1:1000 with BG11 -Mn and 2 µL of these cell suspensions were dropped onto BG11 medium supplemented without (w/o) MnCl_2_ (0 µM), 1x MnCl_2_ (9 µM), or 5x MnCl_2_ (45 µM). Pictures were taken after 5 d growth at 30 °C, 70 µmol photons m^-2^ s^-1^. D) Determination of intracellular Mn concentrations before (0 h) and 4 h after addition of 9 µM MnCl_2_. Cells were precultivated for 5 d in BG11 -Mn and then treated with 9 µM MnCl_2_. Samples were taken before (0 h) and 4 h after the addition of MnCl_2_. To only determine the intracellular Mn pool, periplasmic Mn was eliminated by two washing steps with 5 mM EDTA in 20 mM HEPES buffer (pH 7.5). Data obtained for the WT measurement at 0 h was set to 100 % and subsequent values were normalized to this point. Typical values obtained for the WT at 0 h were 1.4*10^6^ Mn atoms/cell (= 100%). Shown are averages and standard deviations of three biological (with five technical) replicates each. Asterisks indicate significant differences between the reference value and specified knockout mutant according to a Student’s t-test (*: *P* ≤ 0.05; **: *P* ≤ 0.01).

The results support the assumption that Hmx1 and Hmx2 facilitate Mn transport, likely Mn import with high specificity. We additionally examined the intracellular Mn concentrations and found that all mutant lines, whether single or double mutant, accumulated significantly less Mn inside the cell (excluding the periplasmic space) after the addition of 1x MnCl_2_ (Figure 2D). The equal growth and accumulation phenotype of single and double mutants indicated that Hmx1 and Hmx2 need to be both active and probably functionally assemble as heteromers *in vivo*.

### Hmx1 and Hmx2 reside in the plasma membrane

The low Mn sensitive phenotype and the reduced intracellular Mn pools of the mutants pointed toward an import function for Hmx1/2, prompting the investigation of the subcellular localization of the proteins. We generated mutant lines expressing either Hmx1 (*hmx1:cfp*) or Hmx2 (*hmx2:cfp*) fused to a C-terminal cyan fluorescent protein (CFP). The fusion constructs were designed in such a way that they replaced the original gene. That is, *hmx1:cfp* and *hmx2:cfp* were introduced into the native genomic context under control of the respective native promoter. To use these constructs as complementation lines and thus demonstrate that the observed phenotypes are only due to the deletion of the specific gene, we transformed Δ*hmx1* and Δ*hmx2* single-mutant cells. Full segregation of the expression lines demonstrated that the fused version fully displaced the knockout alleles (Figure S2A). In comparison to Δ*hmx1/*Δ*hmx2*, the *hmx1:cfp* and *hmx2:cfp* lines are not sensitive toward low Mn concentrations (Figure S2B). On the one hand this demonstrates that the CFP fusion proteins functionally assembled in the correct native cellular site, and on the other hand that the expression lines could be used to complement the mutants. Confocal fluorescent microscopy (Figure 3) revealed that the CFP signals of both Hmx1 and Hmx2 did not fully overlap with the signal of the chlorophyll autofluorescence, but were rather oriented toward the outside of the cell. This suggests that both Hmx1 and Hmx2 reside in the plasma membrane. Furthermore, the CFP signals had a patchy, dotted appearance and showed local maxima at regions of low chlorophyll fluorescence (e.g., Figure 3B: ROI6 and ROI9; Figure 3D: ROI6 and ROI8), which are typical for PSII biogenesis centers (Heinz et al., 2016).

**Figure 3:**
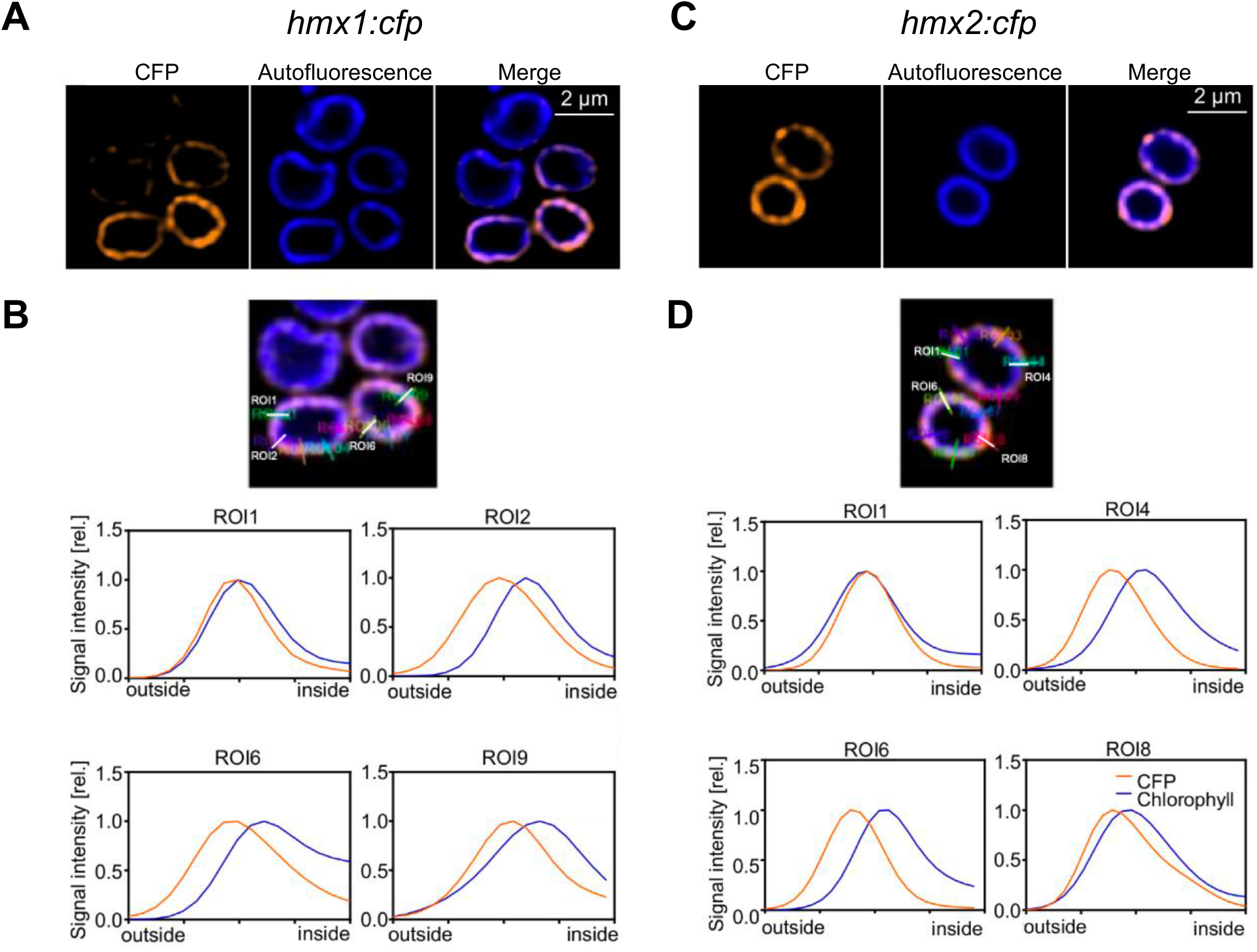
Subcellular localization of Hmx1 and Hmx2. The subcellular localization of Hmx1 and Hmx2 was determined by confocal fluorescence microscopy. CFP was C-terminally fused to Hmx1 and Hmx2. The fusion constructs were introduced into the original *hmx1* and *hmx*2 locus, resulting in the strains *hmx1:cfp* and *hmx2:cfp* and expression under the endogenous promoter. Typical results for *hmx1:cfp* are shown in A and B and for *hmx2:cfp* in C and D. CFP fluorescence is shown in orange, chlorophyll autofluorescence in blue, and a merged image shows both signals at the same time (A and C). Intensities of the signals from the outside to the inside of the cell at various positions of the cell circumference were analyzed as indicated by the regions of interest (ROIs) (B and D). Intensities of the signals along the cell circumference were analyzed as indicated by the ROIs (B and D).

### Double mutants in *hmx2* and *mntC* are not viable under Mn-limiting conditions

Previous work identified MntCAB as the major Mn importer at the plasma membrane under Mn-limiting conditions (Bartsevich & Pakrasi, 1996). Since in those experiments Mn uptake was not fully abolished in a Δ*mntC* mutant, the presence of a second high-affinity transport system was postulated. Hmx1/2 also reside in the plasma membrane and likely facilitate Mn transport. To provide further evidence for their transporter function, we generated a single mutant in *mntC,* Δ*mntC* (Figure 4A) and a double mutant with an additional defect in *hmx2*, Δ*mntC/*Δ*hmx2.* In the Δ*mntC* mutant, zero MntCAB transport activity was observed (Bartsevich & Pakrasi, 1996). After genotype verification (Figure 4B), the mutant lines were studied for their Mn sensitivity (Figure 4C). As expected, Δ*mntC* showed strong growth retardation only on medium lacking Mn. A concentration of 0.5x (4.5 µM) MnCl_2_ in the BG11 medium was sufficient to fully compensate the phenotype. The Δ*hmx2* mutant underperformed under reduced Mn availability conditions (w/o to 2.5x MnCl_2_), but could be rescued by elevated (5x) Mn concentrations. The double mutant mounted the strongest Mn sensitivity phenotype. A MnCl_2_ concentration below 1x was not sufficient to allow Δ*mntC/*Δ*hmx2* to grow. However, 5x MnCl_2_ in the medium fully rescued the phenotype. Likewise, the determination of intracellular Mn demonstrated that Δ*mntC* and Δ*mntC/*Δ*hmx2* contained significantly depleted Mn pools after a 5-day period of Mn limitation (Figure 4D). Furthermore, the addition of 0.5x MnCl_2_ resulted in disturbed intracellular accumulation in all mutant lines, with the most pronounced impairment observed in Δ*mntC/*Δ*hmx2.* The accumulation was significantly reduced even in comparison to both single mutants, Δ*hmx2* and Δ*mntC* (Figure 4D).

**Figure 4:**
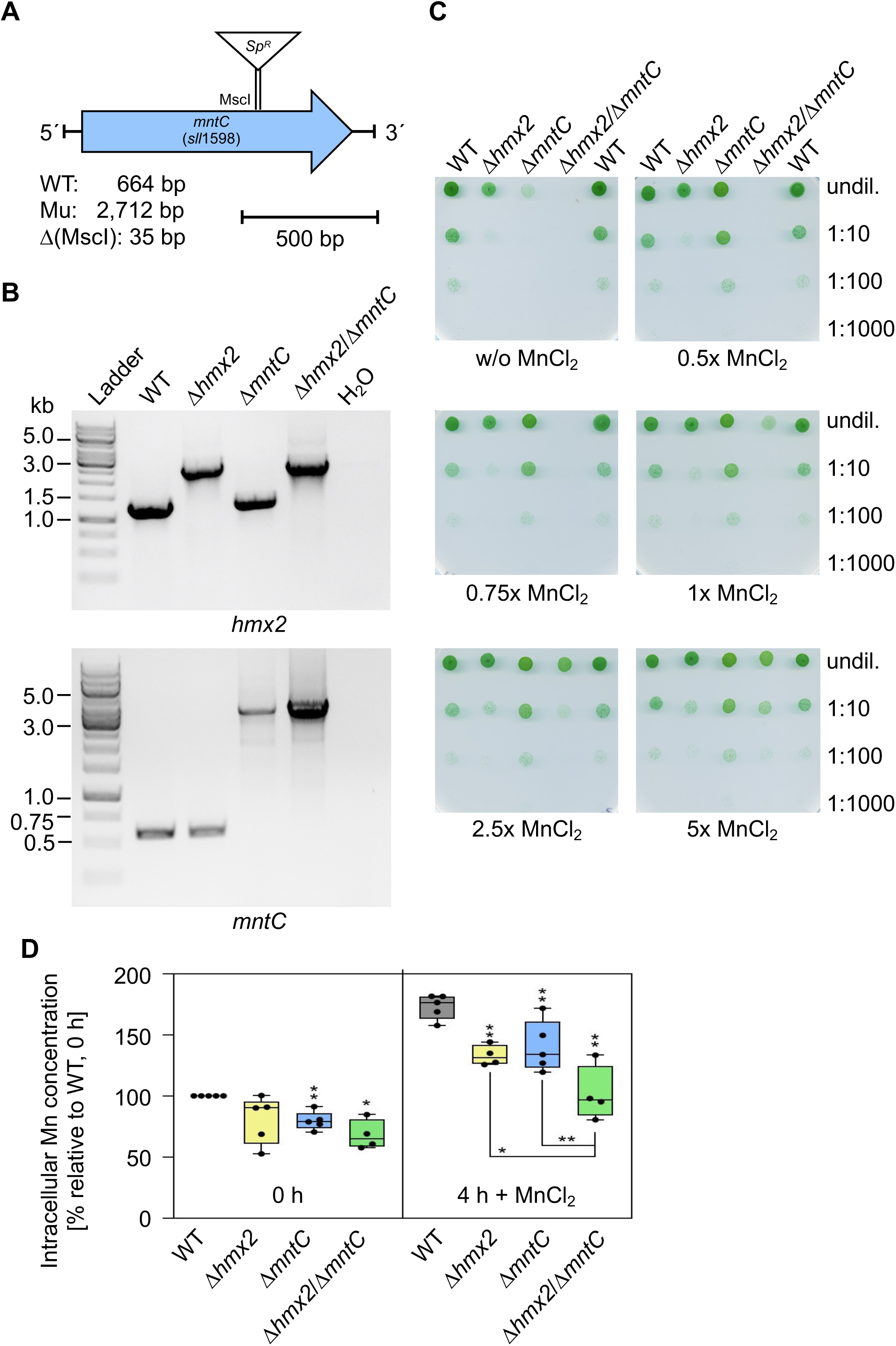
Analysis of Δ*hmx2* and Δ*mntC* mutants. A) Scheme for the generation of a mutant in *mntC* (Δ*mntC*) by deletion (Δ = 35 bp) and insertion of a spectinomycin resistance cassette (Sp^R^). B) Verification of mutations in Δ*hmx2*, Δ*mntC,* and Δ*hmx2/*Δ*mntC* by genotyping. PCR analysis was performed with gDNA and gene specific primers (*hmx2*: primers ME368/ME369, WT = 1,279 bp; Mu = 2,531 bp. *mntC*: primers ME163/ME164, WT = 664 bp; Mu = 2,712 bp). C)Droptest to monitor Mn sensitivity. Cells were washed with BG11-Mn and adjusted to an OD_750_ of 0.25. Cells were diluted 1:10, 1:100, 1:1000 with BG11-Mn and 2 µL of these cell suspensions were dropped onto BG11 medium supplemented without (w/o) MnCl_2_ (0 µM), 0.5x MnCl_2_ (4.5 µM), 0.75x MnCl_2_ (6.75 µM), 1x MnCl_2_ (9 µM), 2.5x MnCl_2_ (22.5 µM), or 5x MnCl_2_ (45 µM). Pictures were taken after 5 d growth at 30 °C, 70 µmol photons m^-2^ s^-1^. D) Determination of intracellular Mn concentrations before (0 h) and 4 h after addition of 4.5 µM MnCl_2_. Cells were precultivated for 5 d in BG11 -Mn and then treated with 4.5 µM MnCl_2_. Samples were taken before (0 h) and 4 h after the addition of MnCl_2_. To exclusively determine the intracellular Mn pool, periplasmic Mn was eliminated by two washing steps with 5 mM EDTA in 20 mM HEPES buffer (pH 7.5). Data obtained for the WT measurement at 0 h was set to 100 %, and subsequent values were normalized to this point. Typical values obtained for the WT at 0 h were 1*10^6^ Mn atoms/cell (= 100%). Shown are averages and standard deviations of 4 or 5 biological with 5 technical replicates each. Asterisks indicate significant differences between referring WT value and specified knockout mutant according to a Student’s t-test (*: *P* ≤ 0.05; **: *P* ≤ 0.01).

### *hmx1* and *hmx2* are constitutively expressed

To test for house-keeping or inducible gene expression, the coefficient of variation was calculated from 46 publicly available data sets in the short read data archive (Wulf et al., 2024) and showed that *hmx1* and *hmx2* belong to the transcripts of low variation, whereas *mntCAB* were highly variable transcripts (Figure 5A). Mn-independent expression of *hmx1* and *hmx2* was additionally supported by analyzing transcriptional profiles of *Synechocystis* WT cells from Mn limitation (Sharon et al., 2014) and Mn excess (Reis et al., 2024) experiments. While transcript abundances of the *mntCAB* operon were elevated under Mn-limiting conditions and reduced under Mn excess conditions, *hmx1* and *hmx2* expression was not significantly changed under either condition (Figure 5B).

**Figure 5:**
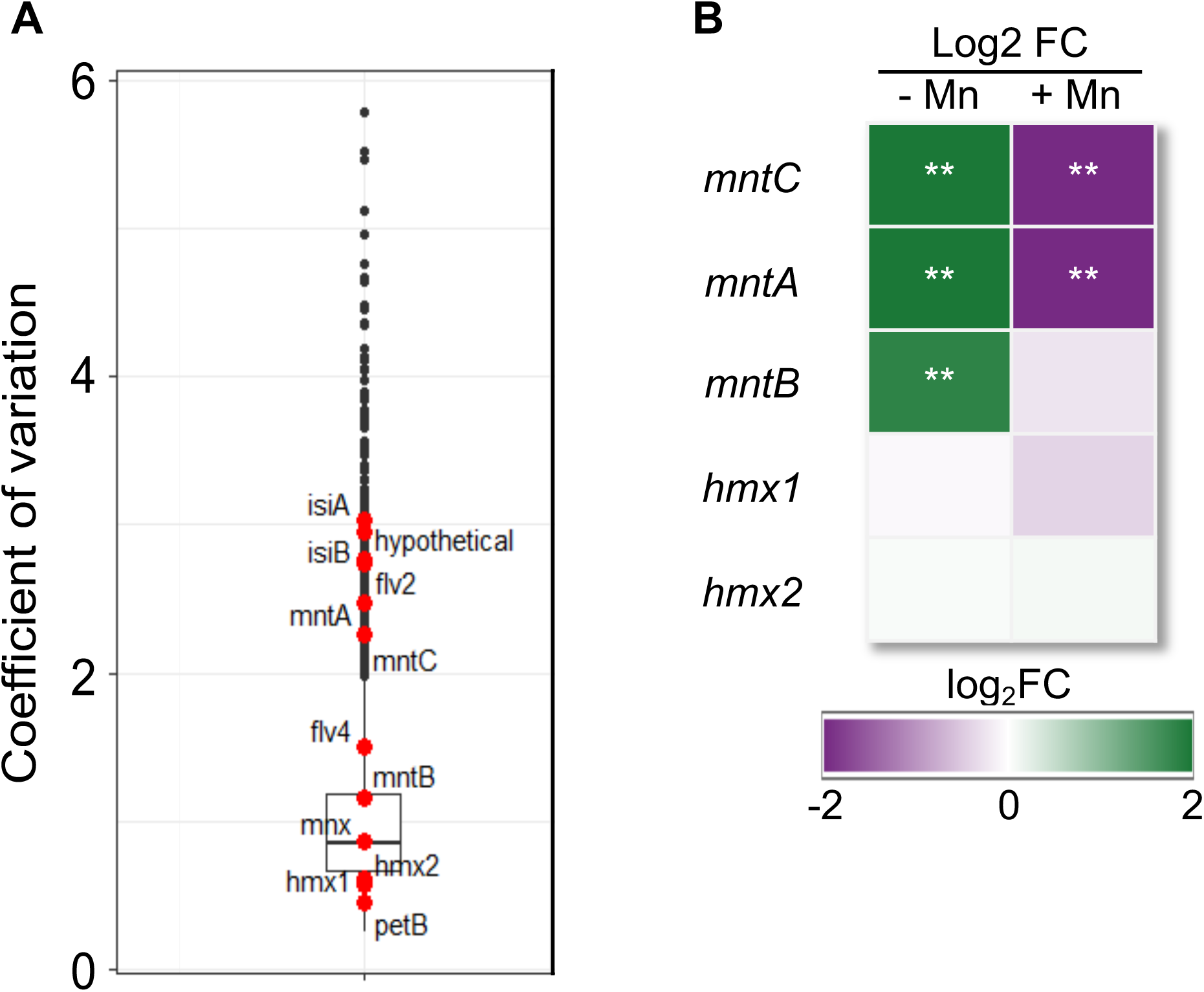
Expression analysis of *hmx1* and *hmx2.* A) Coefficient of variation for *hmx1* and *hmx2.* As representatives for inducible gene expression, *flv2*, *flv4*, *isiA*, *isiB*, *mntA*, *mntB*, and *mntC* were visualized. *petB* is used as a control for a house-keeping gene. B) Mn-dependent transcript accumulation in *Synechocystis* WT cells. Given are log2 fold changes of transcript abundances 48 h after application of Mn limitation (w/o MnCl_2_) conditions (-Mn; Sharon et al., 2014) or 24 h of Mn excess (10x MnCl_2_) conditions (+Mn, Reis et al., 2024) versus Mn control (1x MnCl_2_) conditions. Asterisks indicate significant changes (*q* < 0.01, q-value calculated according to Benjamini-Hochberg).

### Hmx1 and Hmx2 are conserved across and are almost exclusive to cyanobacteria

To determine the evolutionary origin of the investigated Mn transporters, a database of 5,455 bacterial and 212 archaeal complete genomes was screened via diamond BLASTp. Since Mnx, Hmx1, and Hmx2 share substantial sequence identity, additional pairwise global alignments were used to determine the annotation of all hits. The gene distribution suggests a cyanobacterial origin of Hmx1 and Hmx2, since Hmx-type homologs with three TMDs are rarely found outside of cyanobacteria, independent of the chosen e-value cutoff (Figure 6, Figure S3, Table S2). Exceptions are single Hmx1/Hmx2 homologs in *e.g*., *Desulfovibrio desulfuricans* ND132 or *Desulfovibrio magneticus* RS-1. In contrast, Mnx homologs with six TMDs are present in a wider range of Proteobacteria, Actinobacteria, Chlorobi, Clostridia, and one member of the Negativicutes. Eukaryotic genomes appear to encode only Mnx homologs having six TMDs conserved (Figure S4, Table S2). Focusing on the cyanobacterial genomes (Figure 6), it becomes apparent that some strains contain all three homologs (Hmx1, Hmx2, and Mnx; *e.g*., *Synechocystis* sp. PCC 6803), some only the pair of Hmx1 and Hmx2 (*e.g*., *Synechococcus* WH 7803), and a few only one Hmx homolog (*e.g*., *Gloeobacter violaceus* PCC7421). Cyanobacterial genomes hence offer a window into the stages of Hmx/Mnx evolution, ranging from the single half-sized, 3-TMD homolog of the UPF0016 family in an early branching cyanobacterium, such as *Gloeobacter*, via two 3-TMD Hmx homologs in the majority of branching cyanobacteria, to the 6-TMD version of UPF0016 homologs in about half of them.

**Figure 6:**
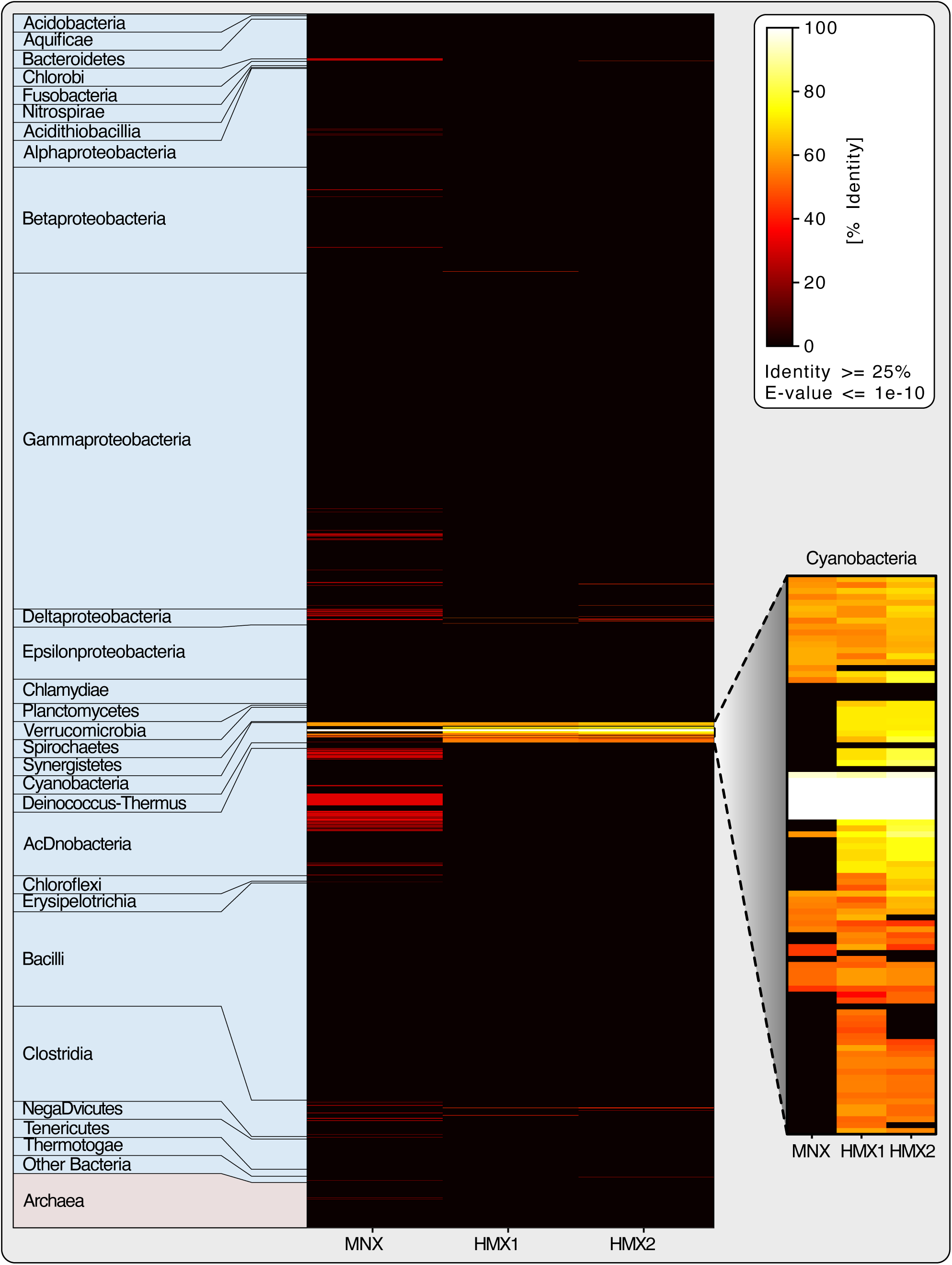
Occurrence of UPF0016 genes in prokaryotic genomes. A database of 5,655 complete prokaryotic genomes was searched for homologs of Mnx, Hmx1, and Hmx2 via DIAMOND. The presence-and-absence pattern was color coded based on sequence identity and only hits with at least 25 % sequence identity and a maximum e-value of 1E-10 were plotted. Occurrence of Mnx, Hmx1, and Hmx2 homologs in cyanobacterial genomes only are shown in the inlet.

## DISCUSSION

Members of the UPF0016 family have recently emerged as important representatives of secondary transporters with Ca^2+^ and, most importantly, Mn^2+^ as substrates (Stribny et al., 2020). Typically, the 6-TMD transporters occur in a wide range of eukaryotic groups (Figure S4) and also cyanobacteria (Figure 6). In eukaryotes the transporters serve in Ca^2+^ and Mn^2+^ uptake into the Golgi apparatus, where Mn-dependent glucosyltransferases are located. A functional defect of the human homolog TMEM165 is linked to cases of congenital disorders of glycosylation, an inheritable disease leading to various pathological symptoms (Foulquier et al., 2012; Stribny et al., 2020).

In photosynthetic eukaryotes, genes encoding 6-TMD members of the UPF0016 family have undergone multiple duplication events (Hoecker et al., 2017). The genome of *Arabidopsis* for instance encodes five UPF0016 proteins (Hoecker et al. 2017). CHLOROPLAST MANGANESE TRANSPORTER 1 (CMT1) and PHOTOSYNTHESIS-AFFECTED MUTANT 71 (PAM71) reside in plastid membranes. CMT1 imports Mn across the plastid inner envelope into the plastid stroma, while PAM71 facilitates uptake into the thylakoid lumen to provide Mn for its incorporation into the oxygen evolving complex of PSII (Schneider et al., 2016; Eisenhut et al., 2018; Zhang et al., 2018). PHOTOSYNTHESIS-AFFECTED MUTANT 71 LIKE 3 (PML3), also known as BIVALENT CATION TRANSPORTER 3 (BICAT3), is *trans*-Golgi localized and plays a critical role in glycosylation reactions under Mn limitation conditions, in cell wall biosynthesis (Yang et al., 2021), and the allocation of Mn between Golgi apparatus and chloroplast (He et al., 2022). PML4 and PML5 localize to the endoplasmic reticulum and likely fine-tune the uptake of Mn (Hoecker et al., 2020). We identified half-sized versions of the UPF0016 proteins, Hmx1 and Hmx2, in the cyanobacterium *Synechocystis* that each contain three instead of six TMDs (Figure 1A).

### Hmx1 and Hmx2 enable constitutive Mn uptake as heteromers at the plasma membrane

The vast majority of cyanobacterial genomes encode two genes encoding 3-TMD proteins, where *hmx1* and *hmx2* are usually arranged in an operon (Figure 1B, Table S3). Since such regulatory systems typically contain genes coding for functionally related partners, we expected both proteins Hmx1 and Hmx2 to assemble as heteromers and function as Mn transporter in the same way as their 6-TMD homologs of the UPF0016 family. Though a direct physiological interaction was not investigated by us, our results support this hypothesis.

We were able to delete the genes in single and double knock-out mutants (Figure 2B). The full genome segregation indicates that Hmx1 and Hmx2 are not essential for survival under standard conditions with 1x (9 µM) MnCl_2_ supplementation. The reduced growth performance at this concentration and even more so in the absence of Mn (Figure 2C), however, clearly demonstrates its important role in Mn uptake. The decreased ability to accumulate Mn inside of cells after a 9 µM MnCl_2_ pulse (Figure 2D) furthermore corroborates the Mn uptake function for Hmx1/2.

The only Mn ABC-type transporter MntCAB that has been described for cyanobacteria until now, serves Mn import at the plasma membrane under Mn limitation (Bartsevich & Pakrasi, 1995; Bartsevich & Pakrasi, 1996). Intriguingly, *mntCAB* knockout lines continued to show Mn uptake when supplemented with micromolar amounts of Mn, indicating the presence of a second, yet unidentified Mn uptake system (Bartsevich & Pakrasi, 1996). Our analysis of double mutants with deletions in both *mntC* and *hmx2* revealed that both transporters assist each other at Mn scarcity, since Δ*mntC/*Δ*hmx2* mutants are not able to survive without any Mn supplementation (Figure 4C), while also showing the lowest levels of intracellular Mn accumulation (Figure 4D). Bartsevich and Pakrasi (1996) furthermore postulated that the additional Mn transport system should be active at different, likely micromolar extracellular Mn conditions. This holds true for Hmx1/2. While the Δ*mntC* mutant shows the strongest growth retardation at 0x MnCl_2_, the phenotype is already fully recued at 0.5x (4.5 µM) MnCl_2_. This result was expected, since expression of *mntCAB* occurs under the control of the ManS/ManR two-component system (Ogawa et al., 2002; Yamaguchi et al., 2002) and is known to be induced only at extracellular Mn concentrations below 1 µM (Yamaguchi et al., 2002; Eisenhut, 2020). Thus, at 4.5 µM MnCl_2_ MntCAB is not expressed and another Mn uptake system must compensate for its absence, which is Hmx1/2. Δ*hmx2* shows impaired growth performance at concentrations up to 5x MnCl_2_ (Figure 2C, Figure 4C) and the Δ*hmx1* and Δ*hmx2* mutants both display Mn sensitivity over a rather broad range of MnCl_2_ concentrations (0 - 9 µM) (Figure 2C). Together with the observation that their gene expression pattern mirrors that of house-keeping genes rather than being inducible (Figure 5A), and hardly reacts to changes in Mn supplementation (Figure 5B), we postulate that Hmx1/2 functions as a constitutive Mn transporter.

It was furthermore proposed (Bartsevich & Pakrasi, 1996) that the unidentified Mn transporter should be highly specific for Mn and our results support this suggestion, too. Besides Mn, only Ca has been demonstrated to be a substrate for some UPF0016 proteins (Demaegd et al., 2013; Colinet et al., 2016; Wang et al. 2016; Frank et al., 2019). For its 6-TMD homolog Mnx, specifically Mn-dependent effects were shown (Brandenburg et al., 2017b; Gandini et al., 2017). Our experiments with medium depleted of Ca or Fe (Figure S1) revealed unaffected growth of Δ*hmx1* and Δ*hmx2* mutants. Though we cannot rule out transport of other untested metals, such as copper or zinc, our results point towards a high substrate specificity of Hmx1/2 for Mn.

Finally, to serve Mn import, the proteins are expected to localize to the plasma membrane. In both lines expressing *hmx1:cfp* or *hmx2:cfp* we observed that the CFP fluorescence did not entirely overlap with the chlorophyll autofluorescence (Figure 3), which would be indicative for a thylakoid membrane localization. The halo-like appearance is different from thylakoid membrane proteins, such as Mnx (Brandenburg et al., 2017b) and rather indicates Hmx1 and Hmx2 residing in the plasma membrane. To minimize experimental artifacts, the CFP fusion constructs were introduced into the geneś native chromosomal sites, so that expression was guided by the native promoters and to rule out potential overexpression artifacts. Furthermore, since the expression lines did not show Mn sensitivity (Figure S2B), we argue that Hmx1:CFP and Hmx2:CFP reside in their designated locations and are functional.

Uptake across the plasma membrane serves Mn delivery to cytoplasmic Mn-requiring proteins and also further passage via the thylakoid membrane transporter Mnx to assist Mn incorporation into PSII. The additional and distinct patchiness of the CFP fluorescence signal is very similar to that observed for CurT (Heinz et al., 2016; Ostermeier et al., 2022). CurT locates to thylakoid convergence zones, which enable contact sites between the plasma and thylakoid membrane. These zones are postulated to house early steps in PSII assembly (Nickelsen & Rengstl, 2013; Heinz et al., 2016). Thus, it is conceivable that Hmx1/2 resides at these contact sites and enables the Mn uptake for efficient provision of the metal cofactor at the site of pD1 translation, PratA-assisted Mn incorporation, and PSII biogenesis. Further studies, however, will be needed to prove this hypothetical function.

Strikingly, the compensation of the Mn sensitive phenotype *of* Δ*hmx2* (Figure 2C) and the viability of the Δ*mntC/*Δ*hmx2* double mutant (Figure 4C) at elevated Mn concentrations suggest the presence of an additional Mn uptake system at the plasma membrane. However, this transporter likely has only a low-affinity for Mn as indicated by the compensation with 5x MnCl_2_ (at minimum). A good candidate for the low-affinity Mn transport system is the Fe(III) ABC-type transporter FutABC (Katoh et al., 2001). Expression of the transporter subunits is slightly enhanced at Mn limitation (Sharon et al., 2014). FutABC might transport Mn as co-substrate by a piggybacking mechanism (Brandenburg et al., 2017b; Eisenhut, 2020).

Hmx1 and Hmx2 are rather small proteins (117 and 92 amino acids, respectively). They both comprise three TMDs and contain the signature ExGD motive, which is suggested to participate in forming the pore of the transporter (Stribny et al., 2020). According to our experimental results, they are only functional if both are present, presumably assembling as heteromers: single and double mutants in *hmx1* and *hmx2* have congruent phenotypes. That is, they show reduced growth on medium with low Mn concentrations (Figure 2C) and have smaller intracellular Mn pools in comparison to the WT (Figure 2D). If homomers were active in Mn transport, single deletions of either *hmx1* or *hmx2* would not have resulted in Mn sensitivity and additive characteristics in the Δ*hmx1/*Δ*hmx2* double mutant would have been expected. Also, *hmx1* and *hmx2* are with 90 % (Table S3) predominantly encoded as operons, which supports a functional interaction of both 3-TMD proteins and single 3-TMD units are too small for building a transporter pore. Thus, we suggest that Hmx1/2 is required to assemble as multimeric heteromers to enable Mn transport. Schneider et al. observed interaction of PAM71, a 6-TMD protein of the UPF0016, with itself (Schneider et al., 2016). Likely, as demonstrated for SWEETs (Xuan et al., 2013) or MFS eukaryote sugar transporters (Abramson et al., 2003), UPF0016 proteins form a functional transporter, if assembling into 12-TMD complexes. Future interaction and structural studies could provide evidence that the 3-TMDs and their conserved ExGD motif, as represented by Hmx1 and Hmx2, in all cases act as the principle building block.

In summary, through the characterization of Hmx1/2 we propose to having identified the so far only biochemically evidenced Mn transporter of cyanobacteria. It allows constitutive Mn uptake and in collaboration with the MntCAB system ensures proper Mn provision under both Mn-limiting and sufficient conditions. A model with Hmx1/2 integrated into the Mn homeostasis network is provided in Figure 7. Like chloroplasts of the green lineage (Chloroplastida), also cyanobacteria employ two transporters of the UPF0016 family for sequential uptake of Mn. A comparable example is the occurrence of Cu-specific P-type ATPases in both cyanobacteria (CtaA and PacS) (Kanamaru et al., 1994; Phung et al., 1994; Tottey et al., 2001) and plants (PAA1 and PAA2) (Abdel-Ghany et al., 2005). In both cases the homologs act in tandem to transport Cu^+^-ions via the plasma membrane/chloroplast envelope and the thylakoid membrane.

**Figure 7:**
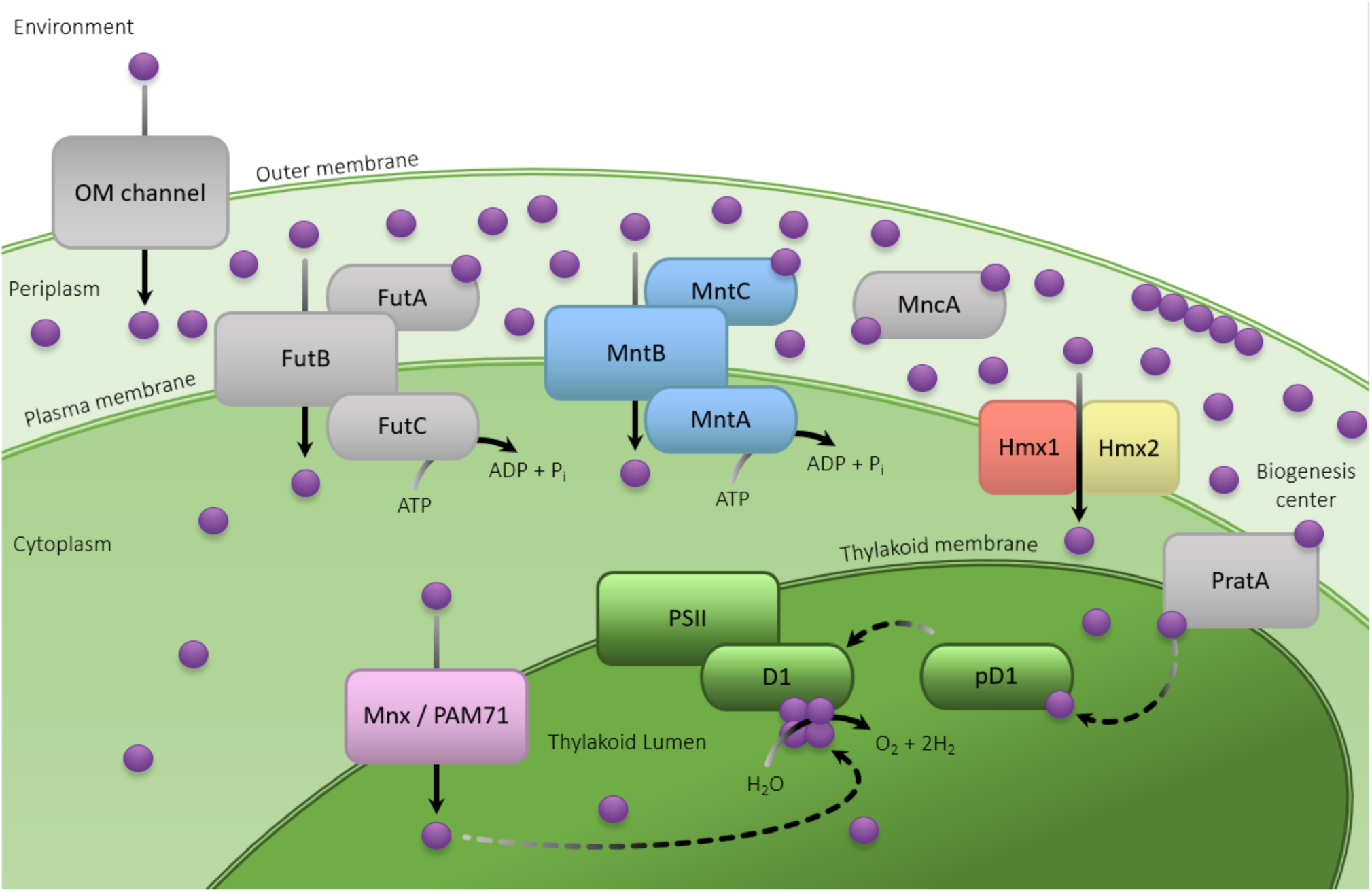
Biological role of Hmx1 and Hmx2 in cellular Mn homeostasis of cyanobacteria. For *Synechocystis*, two distinct Mn pools were observed: up to 80% of the cellular content was found to accumulate in the periplasm, while only 20% reside inside the intracellular space (Keren et al., 2002). In the periplasmic space, Mn is most likely loosely bound to the outer membrane by the negative membrane potential or bound to soluble Mn-binding proteins, such as MncA (Keren et al., 2002; Tottey et al., 2008). Inside the cell, roughly 80% of the Mn is associated with the Mn-cluster of the OEC. This major sink of Mn is assumed to being supplied via two routes. i) The tetratricopeptide repeat protein PratA delivers Mn from the periplasm directly to the precursor of the D1 (pD1) reaction center protein in the biogenesis centers of the cell (Stengel et al., 2012). ii) The manganese exporter Mnx (Brandenburg et al., 2017b), also known as SynPAM71 (Gandini et al., 2017) transports Mn from the cytoplasm into the thylakoid lumen and thus serves as an alternative delivery option. Besides the ABC-type transporter MntCAB, which is only expressed under Mn-limiting conditions (Bartsevich & Pakrasi, 1996), the here studied UPF0016 proteins Hmx1 and Hmx2 serve Mn import at the plasma membrane and biogenesis centers. They form heteromers and are constitutively active. The ferric Fe importer FutABC likely has a low-affinity Mn uptake activity.

### Cyanobacterial Hmx1 and Hmx2 represent the ancestral form of UPF0016 proteins

Pairs of Hmx proteins are almost exclusively encoded in cyanobacterial genomes (Figure 6). Isolated *hmx* genes, so called singletons, are found in some cyanobacteria, including *Gloeobacter*, but also in other microorganisms (Demaegd et al., 2013). *hmx1* and *hmx2* likely evolved by duplication of a single *hmx* gene within the cyanobacterial clade and were passed on via horizontal gene transfer, also into non-cyanobacterial prokaryotes. Even screening with a fairly low e-value (Figure S3), resulted in the detection of Hmx homologs with three TMDs (also as singletons) only in a very few cases outside of the cyanobacterial group (Figure 6, Table S2). Cyanobacteria, however, frequently encode the fusion form (UPF0016 proteins) with six TMDs. In contrast to the 3-TMD proteins, this 6-TMD type is also observed in several other bacteria, such as Gammaproteobacteria and the MneA of *e.g., Vibrio cholerae* (Fisher et al., 2016), Deltaproteobacteria, or Actinobacteria (Figure 6). An ancestral origin of UPF0016 proteins and subsequential loss in the vast majority of lineages cannot be ruled out, but parsimony suggests a cyanobacterial origin of Hmx proteins and its duplication accompanied by a fusion event to produce 6-TMD proteins of the Mnx-type, which was then horizontally transferred in some cases (Table S2). Fusion forms of proteins, such as Mnx, have a higher probability to be retained after horizontal gene transfer. Since our experiments with the knockout mutants in *hmx1* and/or *hmx2* demonstrated that both proteins are essential for the function as Mn transporter, Hmx proteins only add an advantage, if both proteins are transferred. Thus, their individual probability of being retained after lateral gene transfer is rather low. It is conceivable that Hmx1 and Hmx2 represent the ancestral progenitor of the fused 6-TMD form of UPF0016 proteins, which was – through endosymbiosis – carried into the eukaryotic tree of life (Hoecker et al., 2021). This is a good time to remember that eukaryote genomes only contain 6-TMD versions of UPF0016 members (Figure S4).

The genomic occurrence of a *hmx* singleton in *Gloeobacter violaceus* allows to speculate about a coevolutionary scenario of the UPF0016 Mn transporters with the internalization of the OEC. *Gloeobacter violaceus* is an early-branching cyanobacterium that lacks thylakoid membranes. Its components of the oxygenic photosynthesis apparatus are located in the plasma membrane and the OEC resides in the periplasm (Raven & Sánchez-Baracaldo, 2021). Accordingly, Mn uptake via the plasma membrane is only necessary to provide Mn for intracellular Mn-dependent enzymes, not for the incorporation into the OEC. For this purpose, the single Hmx protein in *Gloeobacter violaceus,* which possibly functions as homomer, might be sufficient. However, with the emergence of thylakoid-forming cyanobacteria, and the consequent internalization of the OEC into the thylakoid system, more efficient Mn uptake became indispensable. It is conceivable that the duplication of a singleton *hmx* gene into *hmx1* and *hmx2* benefits the number of Mn importer that can be present (gene dosage effect), which assures a sufficient and constitutive Mn uptake at the plasma membrane and at the PSII biogenesis centers. The additional fusion form Mnx resides in the thylakoid membrane and serves Mn import into the thylakoid lumen to assist Mn provision to the OEC. This sequential uptake system specifically for Mn was endosymbiontically passed on to plastid containing eukaryotes.

In conclusion, cyanobacteria employ all organizational structures of UPF0016 proteins, from basic 3-TMD units to fused 6-TMD proteins. Hence, cyanobacteria are the most suitable model to study the evolution and function of this important class of transport proteins. The Mn transport function of Hmx1/2 highlights this as fundamental ancient feature of the UPF0016 family. We suggest that Hmx1/2 coevolved as an essential consequence of the cellular internalization of the OEC (reviewed in Martin et al., 2018) at the basis of cyanobacteria performing oxygenic photosynthesis.

## AUTHOR CONTRIBUTIONS

MR, FB, ME designed the research. MR, FB, MK, SF, AB, SM, and ME performed the research. All authors contributed to data analysis and discussion. MR, FB, MK, SBG, and ME wrote the paper. All authors have read and agreed to this version of the manuscript.

## ACKNOWLEDGEMENTS

We thank Stefanie Weidtkamp-Peters and Thomas Zobel (Center of Advanced Imaging, Heinrich-Heine University Düsseldorf) for excellent technical assistance. We thank Jörg Nickelsen and his team (LMU Munich) for providing us with the codon optimized CFP vector. We acknowledge Lisa M. Heihoff for computational support. This work was funded by the Deutsche Forschungsgemeinschaft (DFG) through the grant EI 945/3-2 and SFB1535 - Project ID 458090666 to ME. SBG is grateful for DFG support through MAdLand SPP2237 (440043394) and the CRC1208 (267205415). We acknowledge support for the publication costs by the Open Access Publication Fund of Bielefeld University and the DFG.

## DATA AVAILABILITY STATEMENT

All relevant data can be found in the manuscript or in the Supporting Information.

## SUPPORTING INFORMATION

Table S1: List of oligonucleotides used in this study

Table S2: Occurrence of Mnx, Hmx1, and Hmx2 homologs across bacteria, archaea, and eukaryotes

Table S3: Occurrence and organization of *UPF0016* genes in cyanobacterial genomes

Figure S1: Growth of mutant strains on medium depleted of Ca or Fe

Figure S2: Examination of *hmx1:cfp* and *hmx2:cfp* expression lines

Figure S3: Occurrence of UPF0016 genes in prokaryotic genomes across varying e-value cut-offs

Figure S4: Occurrence of UPF0016 genes in eukaryotic genomes with cut-off e-value ≦ 1E-10

