## Supporting Information for "Hemi Manganese Exchangers 1 and 2 enable manganese import at the plasma membrane in cyanobacteria"

**Table S1:** List of oligonucleotides used in this study. Restriction sites are in small letters

| **Name** | **Sequence (5'→3')** | **Experiment** |
|---|---|---|
| FB69 | CTATTTATTCTCAGCAATATCGCCC | Generation of Δ*hmx1* |
| FB70 | AACAGCACTTTACCTTCAAAGTCC | Generation of Δ*hmx1* |
| ME368 | GGTGGGAAAACTACGTACCT | Generation of Δ*hmx2* |
| ME369 | CATCGGGCTAAGGCTTTACT | Generation of Δ*hmx2* |
| ME163 | AGGCATCGAACCGATTCC | Generation of Δ*mntC* |
| ME164 | TCATTGCTGGGCATTGGT | Generation of Δ*mntC* |
| FB105 | ctcgagCACCCTGGGAGTTTTCACCA | Generation of *hmx1:cfp* |
| FB106 | gctagcGGCGCTGACCACGTCCC | Generation of *hmx1:cfp* |
| FB107 | gaattcGCTCAGCACCACAAACATTCC | Generation of *hmx1:cfp* |
| FB108 | gaattcTGCGGTTAAAAACTTAAATAGGGG | Generation of *hmx1:cfp* |
| FB109 | ctcgagATTTGATTACCCTGCCTTTATTGG | Generation of *hmx2:cfp* |
| FB110 | gctagcATCTTCTTGGTTAGGCCAAAGTAA | Generation of *hmx2:cfp* |
| FB111 | gaattcATCACCGATCGCAATTTTTCC | Generation of *hmx2:cfp* |
| FB112 | gaattcAGTGTGTTTGTGGCCCCAGT | Generation of *hmx2:cfp* |
| FB114 | CGCTCCTGAAAAAAGGGGA | Verification of CFP-mutants |

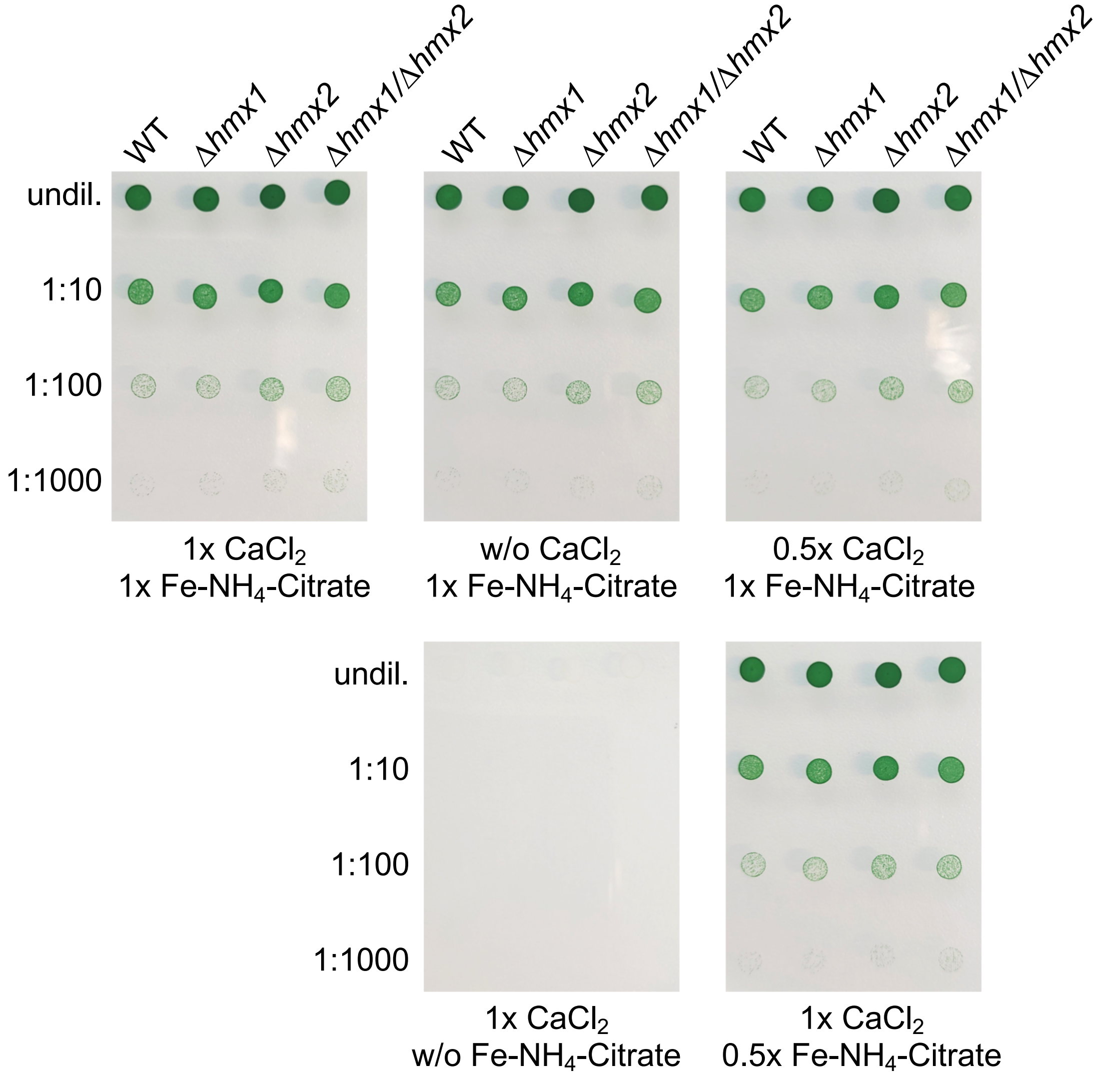


**Figure S1**: Growth of mutant strains on medium depleted of Ca or Fe. Cells were washed with BG11-Ca and -Fe and adjusted to an $OD_{750}$ of 0.25. Cells were diluted 1:10, 1:100, 1:1000 with BG11-Ca/Fe and 2 µL of these cell suspensions were dropped onto BG11 (1x $MnCl_2$) medium supplemented in different combinations of without (w/o) $CaCl_2$ (0 µM) or Fe-$NH_4$-Citrate (0 µg $mL^{-1}$), 0.5x $CaCl_2$ (120 µM) or 0.5x Fe-$NH_4$-Citrate (11.5 µM), and 1x $CaCl_2$ (240 µM) or 1x Fe-$NH_4$-Citrate (23 µM). Pictures were taken after 5 d growth at 30 °C, 70 µmol photons $m^{-2}$.

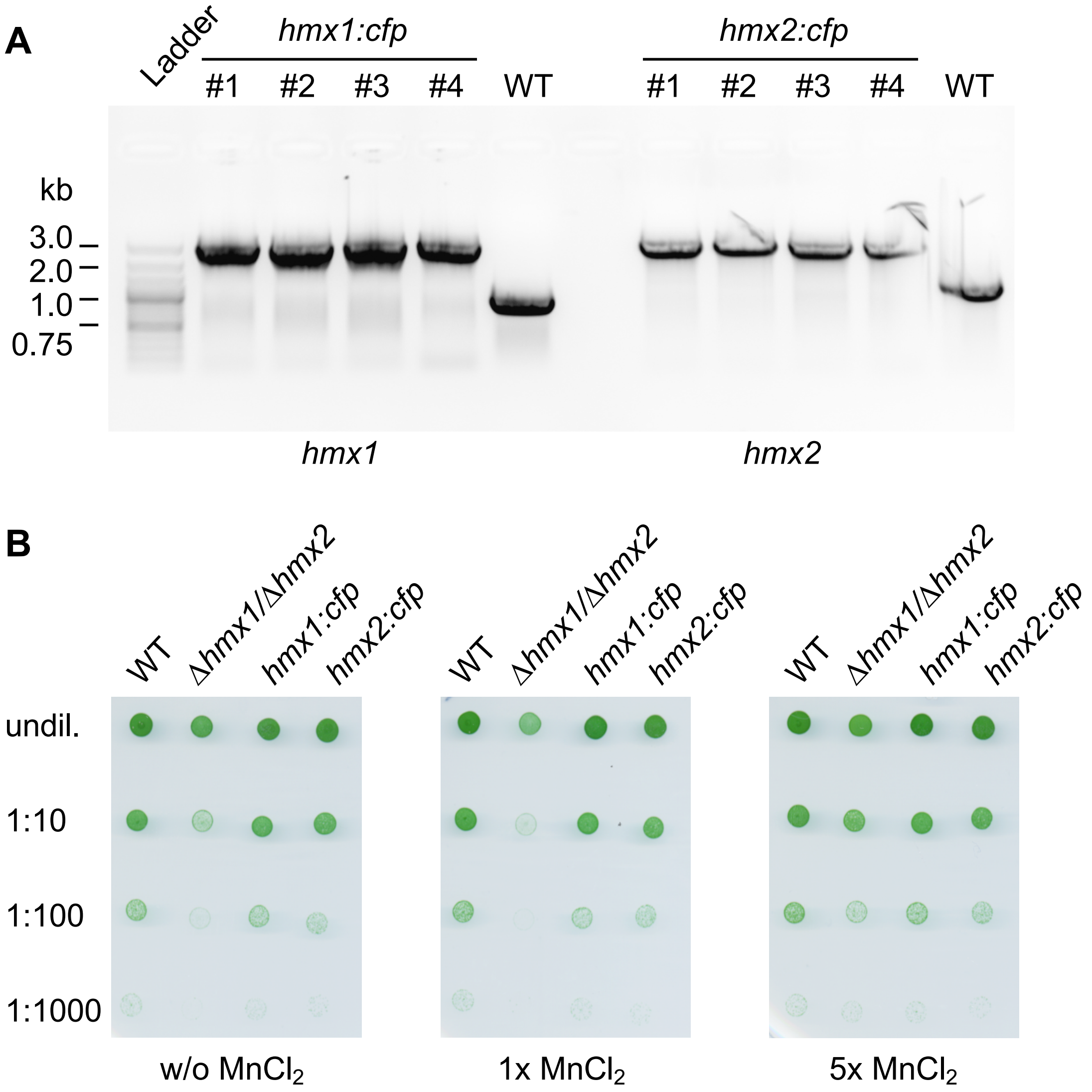


**Figure S2**: Examination of *hmx1:cfp* and *hmx2:cfp* lines. A) Verification of *hmx1::cfp* and *hmx2:cfp* lines by genotyping. PCR analysis was performed with gDNA from four independent clones (#1 - #4) each for *hmx1:cfp* and *hmx2:cfp,* and WT and gene specific primers *(hmx1*: primers FB69/FB70, WT = 954 bp, *hmx1:cfp* = 2,790 bp; *hmx2*: primers ME368/ME369, WT = 1,279 bp, *hmx2:cfp* = 3,115 bp). B) Droptest to monitor Mn sensitivity. Cells were washed with BG11-Mn and adjusted to an $OD_{750}$ of 0.25. Cells were diluted 1:10, 1:100, 1:1000 with BG11-Mn and 2 µL of these cell suspensions were dropped onto BG11 medium supplemented without (w/o) $MnCl_2$ (0 µM), 1x $MnCl_2$ (9 µM), or 5x $MnCl_2$ (45 µM). Pictures were taken after 5 d growth at 30 °C, 70 µmol photons $m^{-2}$.

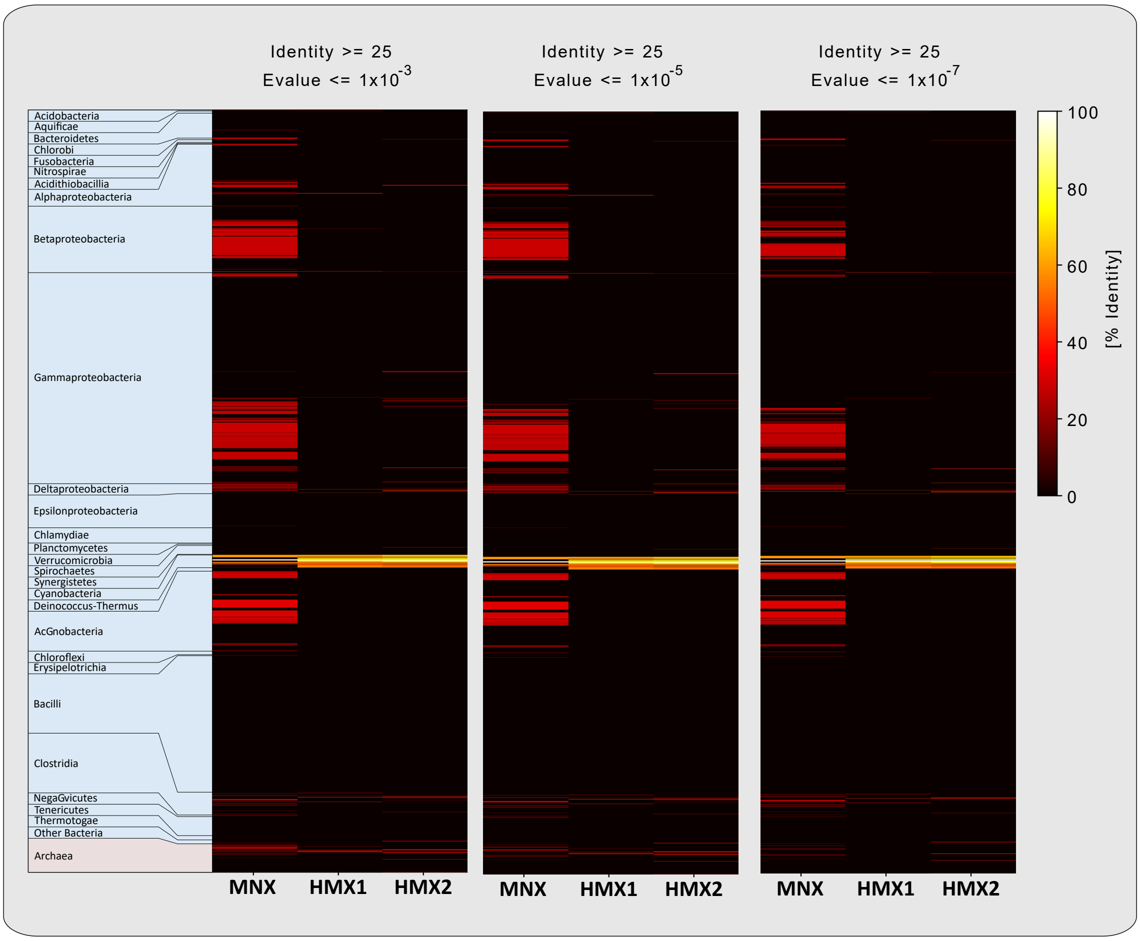


**Figure S3:** Occurrence of UPF0016 genes in prokaryotic genomes with cut-off e-value ≦ 1E-3, e-value ≦ 1E-5, and e-value ≦ 1E-7. The pairwise local identities of each genomes best hit and each query sequence was color coded.

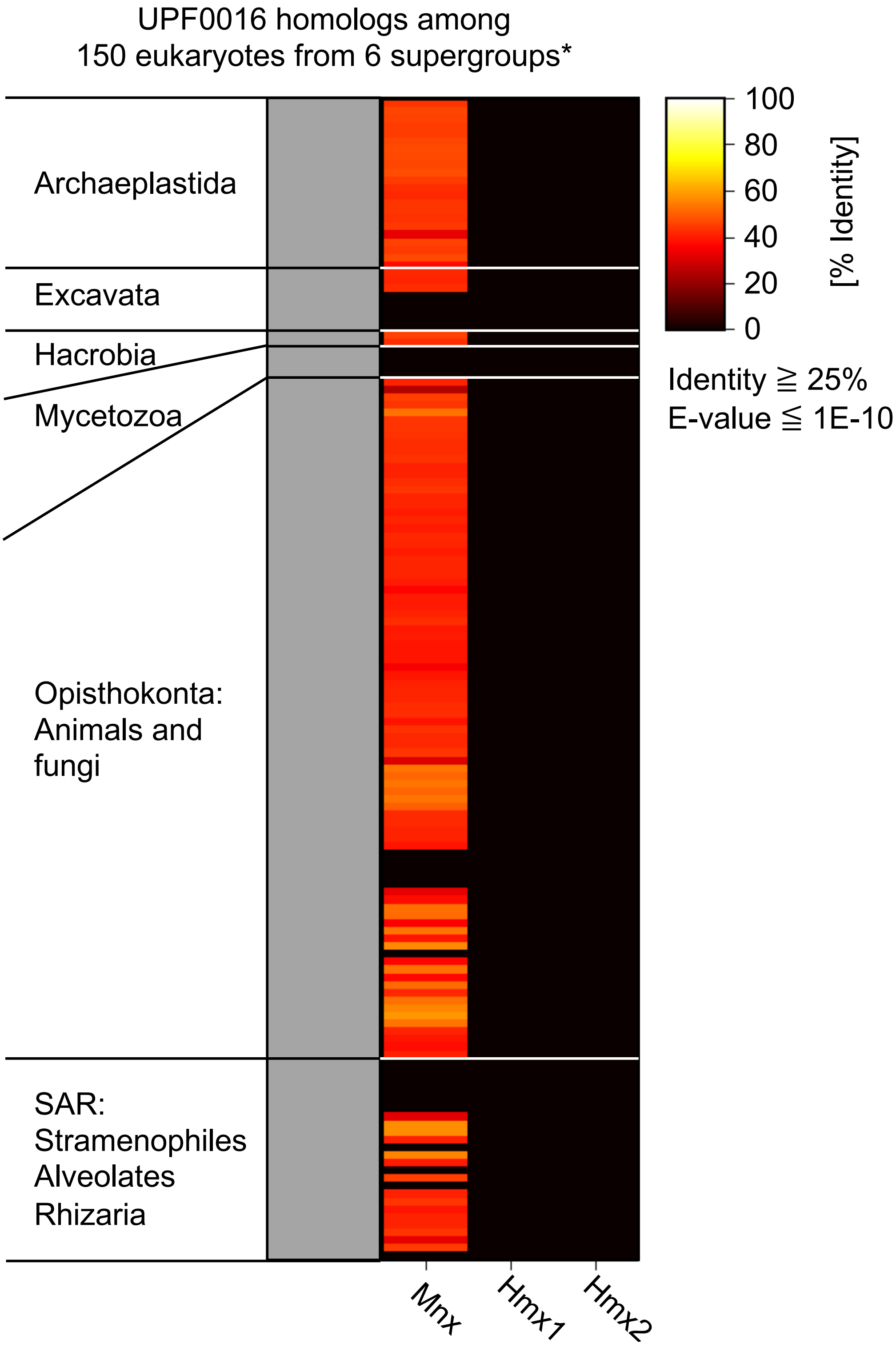


**Figure S4:** Occurrence of UPF0016 genes among 150 eukaryotic genomes with cut-off e-value ≦ 1E-10. The pairwise local identities of each genomes best hit and each query sequence was color coded.
